# On the neural origin of hexadirectional fMRI modulations

**DOI:** 10.64898/2026.09.16.752112

**Authors:** Siyuan Mei, Virginia L. Flanagin, Martin B. Stemmler, Andreas V.M. Herz

## Abstract

When human beings navigate through an environment, a conceptual space, memories or even imagined spaces, fMRI signals in select brain regions vary with movement direction, alternating in strength with a periodicity of 60 degrees. At the cellular level, certain neurons called grid-cells form a hexagonal lattice representation of physical space. Given that the axes of the lattices align across neurons, the concerted directional and spatial firing of populations of these neurons may form the basis of the hexadirectional modulation of the fMRI signal. However, the exact link between the two remains unclear. Multiple mechanisms have been proposed that could couple neuronal activity to the fMRI signal. These include (1) preferential neuronal discharge when the direction of motion coincides with the lattice axes, (2) repetition suppression when multiple fields are traversed in succession, or alternatively, (3) a nonlinear transduction step. Here, we propose targeted experiments that can distinguish these mechanisms, for which we assess the expected effect sizes via simulations. In particular, by introducing calibrated pauses between linear path segments, it is possible to differentiate the repetition-suppression hypothesis from the nonlinear transduction model. By additionally varying the subject’s movement speed and path lengths, the proposed protocol can distinguish conjunctive neuronal tuning — in which neurons respond to both speed and location or direction and location — from the other hypotheses. Crucially, if either the repetition-suppression or the nonlinear-transduction hypothesis holds, then the scale of the lattices will play a role. According to our analysis, hexadirectional signals should emerge only when a linear path segment is longer than half of the spatial lattice period. As grid cells with different scales are grouped into distinct modules along the dorso-ventral axis of the entorhinal cortex, this may make it possible to non-invasively measure the hierarchy of grid scales in the human brain.

## Introduction

Grid cells, first discovered in the rodent entorhinal cortex [1], exhibit spatial firing patterns that form hexagonal lattices covering the explored environment (Fig. 1A). These cells are considered fundamental to navigation. In healthy human beings, single-cell recordings are hardly feasible. However, grid-cell activity has been recorded in epilepsy patients (e.g., [2]), although more often it is inferred instead from hexadirectional fMRI signals [3–20]. These signals were discovered during human navigation experiments in virtual environments [20]. Participants navigate from a random starting point toward a location recalled from memory. During such movements, fMRI signals from the entorhinal cortex are modulated by movement direction *θ* such that a cos(6*θ*)-like pattern emerges, reflecting a six-fold symmetry [20]. While it is tempting to directly attribute this phenomenon to the hexagonal grid-cell activity, a simple population average of the latter produces a uniform, rather than directionally modulated activity [21, 22] (see Fig. 1B).

**Fig 1.**
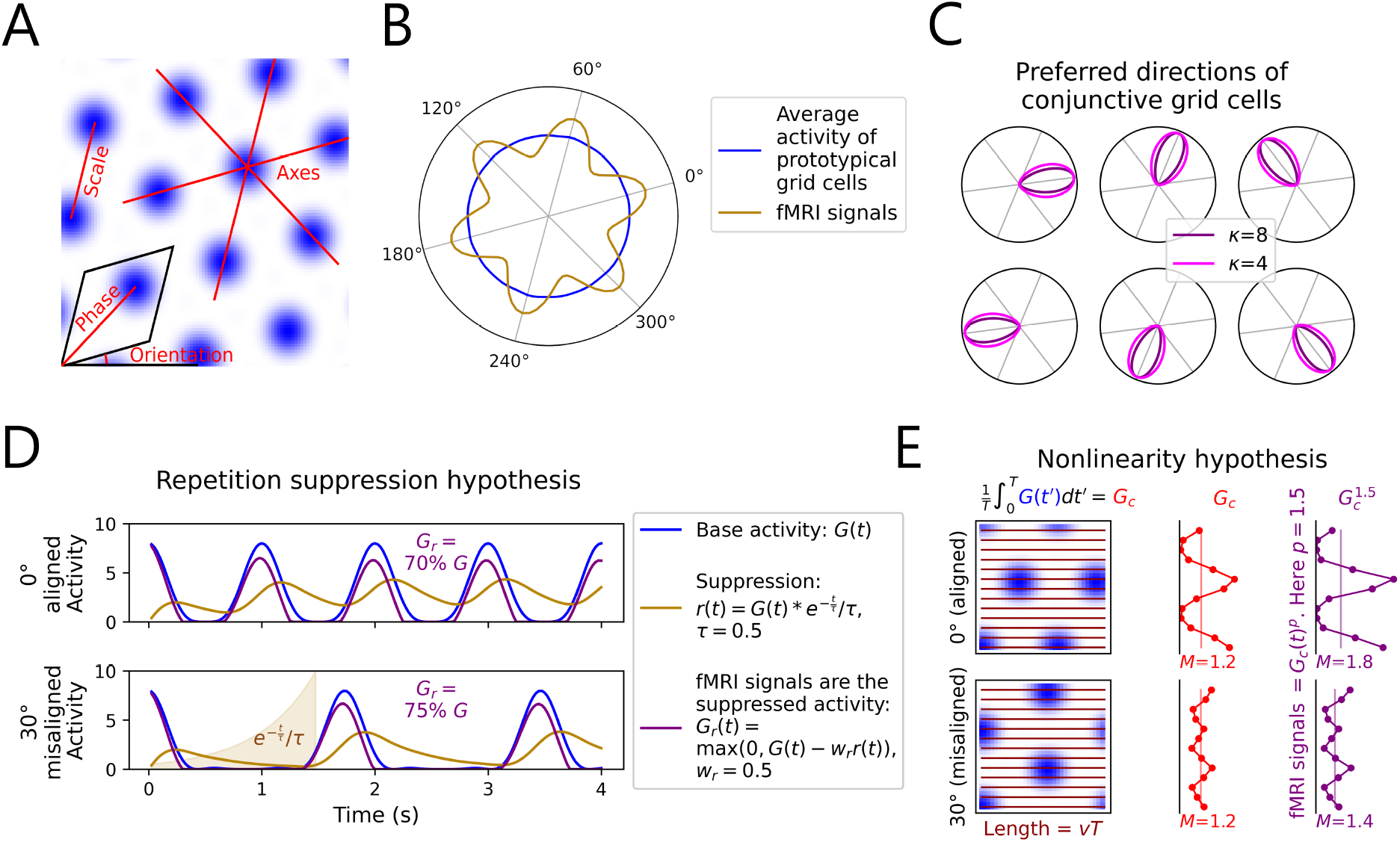
Properties of grid cells and alternative hypotheses for the origin of hexadirectional fMRI signals. **A**. Definition of grid axes, scale (*λ*), orientation, and spatial phase. **B**. The average activity (blue) of grid cells is uniform across movement directions, as seen in simulations. However, hexadirectional fMRI signals (brown) have been observed in several studies that are thought to originate from grid cells. **C**. If grid cells show conjunctive tuning to movement direction or velocity (the first two hypotheses), here modeled by von Mises functions with concentration parameter *κ*, each conjunctive cell should have a single (or multiple) preferred direction(s), with preferred directions across the cell population being periodic in 60°, for hexadirectional fMRI signals to arise. The axes with the strongest responses will have a fixed offset (possibly zero) to the grid axes. **D**. Under the repetition-suppression hypothesis, movement aligned with the grid leads to more frequent re-encounters with the fields of the same grid cell, resulting in stronger suppression *r*(*t*) due to firing-rate adaptation. As a consequence, hexadirectional modulation is observed. See main text for technical details. **E**. Under the nonlinearity hypothesis, the short-time–averaged grid cell activity (*G*_*c*_) exhibits larger spatial variability in the aligned direction, although the average activity *M* is the same across directions. After applying a nonlinear transformation 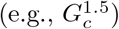, the difference in variability gives rise to the hexadirectional fMRI signature.

To resolve the link between physiology and fMRI, four alternative hypotheses have been proposed. The first two hypotheses rely on grid cells that exhibit conjunctive selectivity to movement direction or velocity [20, 21]. Indeed, in the original study on hexadirectional signals [20], grid cells were found in rodents that were conjunctively tuned to the grid-axis directions (Fig. 1C - **Conjunctive direction hypothesis**). The hypothesis proposed there, was that these cells also exist in human beings and are responsible for the hexadirectional fMRI signals. However, recent analysis [22] on a larger rodent dataset [23] failed to confirm the necessary alignment between preferred directions and grid axes to support this claim. Additionally, given that grid-cell activity persists at rest, the disappearance of the hexadirectional fMRI signal during slow movement [7, 20] speaks against the direction-selective-cell hypothesis. Instead, conjunctive velocity-selective grid cells [24], could solve this discrepancy (**Conjunctive speed hypothesis**). To date, however, no conclusive evidence has been reported that human beings have the conjunctive grid cell properties required for this hypothesis.

The third hypothesis, known as the repetition-suppression mechanism [20, 21], rests on the assumption that grid cells reduce their responses when activated repeatedly in rapid succession (**Repetition-suppression hypothesis**). Moving along the grid axes means that subjects traverse the grid fields of the same cell more frequently than if they move in other directions (Fig. 1D). So firing-rate adaptation should strongly suppress activity in directions aligned with the grid axes, thereby producing hexadirectional fMRI signals. Tentative empirical support for axial modulation of firing-rate adaptation has been found in a recent study [25]. Finally, the **Nonlinearity hypothesis** posits that a nonlinear transformation from single-cell activity to fMRI signal generates the hexadirectional modulation [22]. To obtain this result, Almog et al. first averaged the neuronal activity over a short time window and then raised it to a power greater than unity (Fig. 1E). The nonlinearity is not restricted to power functions and might arise from known nonlinearities in neurovascular coupling [26–29], although direct evidence is missing. Just like the repetition-suppression model, the nonlinearity hypothesis demands that the cells’ firing patterns be grid-like in nature. The two conjunctive hypotheses do invoke intrinsically periodic tuning to direction or velocity, but do not require grid fields—if grid cells contribute, it is because they are conjunctively tuned.

Given the four distinct theories for the origin of the fMRI signal’s hexadirectional modulation, we perform a theoretical analysis to identify and optimize three experimental protocols that will distinguish the four hypotheses. These protocols are exemplified for the case of spatial navigation, involving changes in movement speed, the length of linear path-segments and pause duration between trajectory segments. The protocols can be easily realized in existing navigational paradigms (e.g., [7, 20]); however, one can also envision these manipulations in non-spatial domains.

## Materials and methods

### Simulation procedures

Given the periodic, hexagonal nature of grid-cell activity, we first transformed Cartesian coordinates into a hexagonal coordinate system defined by two axes, *θ* and *ϕ*, separated by 60° and periodic over 2*π*. The spatial scale was then normalized such that a single grid scale *λ* corresponds to 2*π* in this hexagonal space.

Due to this periodicity, the representation was constrained to *θ, ϕ∈*[0, 2*π*), hereafter referred to as the phase (*θ, ϕ*).

For the star-like trajectory simulations, we modeled fMRI signals by averaging over a population of 1,296 grid cells (36*×*36 cells). While these cells shared the same grid scale and orientation, their phases were randomized to reflect biological heterogeneity. Our analysis spanned 100 independent virtual agents across multiple parameter combinations. Each virtual agent’s data was derived from 100 distinct blocks of simulations, each starting from a unique phase, with 36 uniformly distributed movement directions evaluated per block.

For the polar insets in Figures 2C and 2D, we employed the same parameter sets as for the star-like trajectory simulations, with the following modifications to optimize computational efficiency. We reduced the population to 25 grid cells (5*×*5 cells with evenly distributed phases) and conducted only four blocks, each initiated from a unique, uniformly spaced starting phase. These insets represent data from a single virtual agent, with activity tested across 360 equally distributed directions.

**Fig 2.**
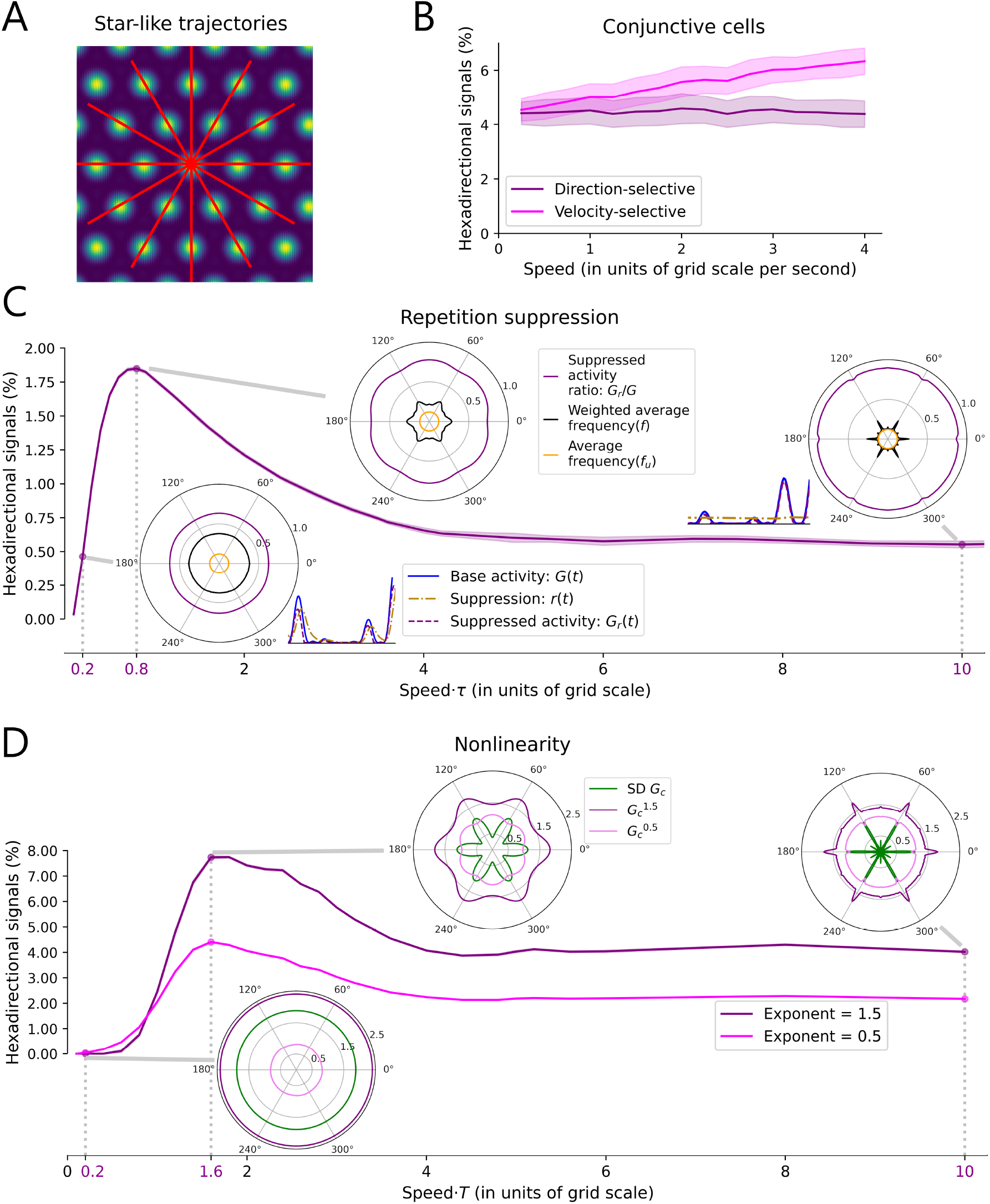
Dependence of hexadirectional signals on movement speed for star-like trajectories. **A**. Example trajectories overlaid on a grid lattice. Every second trajectory overlaps with the lattice. **B**. The hexadirectional signals remain invariant to speed under the conjunctive direction-selective-cell hypothesis, but increase with speed under the velocity-selective-cell hypothesis. The effect of speed under the velocity-selective-cell hypothesis serves as an illustrative example: the modulation can be zero during stationary periods, and the increase can be nonlinear. **C,D**. The hexadirectional signals’ amplitude rises to a single peak as speed increases under both the repetition-suppression hypothesis (**C**) and the nonlinearity hypothesis (**D**). Polar plots illustrate averages of these variables across directions. Insets next to polar plots in (**C**) show examples of activity dynamics under the repetition-suppression hypothesis. SD: standard deviation. Frequency: frequency of encountering the same cell.

Piecewise linear trajectories were simulated using the same protocol as the star-like trajectories, but with a reduced population of 25 grid cells and four blocks. This streamlined configuration was validated by a pilot simulation. To enhance directional detail, we increased the sampling frequency to 60 directions.

### Implementation of grid-cell activity and the four hypotheses

The baseline activity of grid cell *i*, denoted as *G*_*i*_(*t*), was modeled as the product of three cosine waves [21]:

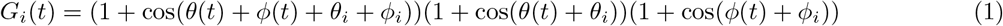

Here, *θ*_*i*_ and *ϕ*_*i*_ represent the spatial phase of grid cell *i* within the hexagonal coordinate system.

For the conjunctive direction-selective grid cells, our simulation followed [21]. Approximately 33% of the grid-cell population was randomly designated as conjunctive. Each conjunctive cell was assigned a preferred direction *ψ*_*i*_ chosen from the six grid-axis directions (multiples of 60°). The directional tuning was modeled using a von Mises distribution, such that the activity *G*_*di*_(*t*) of a conjunctive cell *i* was defined as:

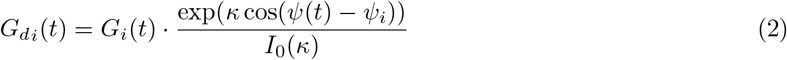

where *κ* represents the concentration parameter (with higher values indicating narrower tuning), *ψ*(*t*) is the current trajectory heading, and *I*_0_(*κ*) is the modified Bessel function of the first kind of order zero. The aggregate fMRI signal was then calculated as the mean activity across both conjunctive and non-conjunctive subpopulations.

For the conjunctive velocity-selective grid cells, we multiplied the aggregate fMRI signals from the direction-selective-cell hypothesis by 1 + *v*/100 when the movement direction was aligned with the grid axis. Here, *v* is the movement speed.

To model the repetition-suppression hypothesis, the suppression of grid cell *i*, denoted as *r*_*i*_(*t*), was defined as a temporal convolution of the baseline activity *G*_*i*_(*t*) with an exponential decay kernel [21]:

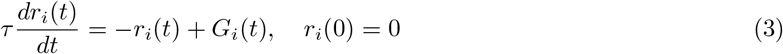

The suppressed activity, *G*_*ri*_(*t*), was then calculated by subtracting the *w*_*r*_-scaled suppression term from the baseline activity, followed by a rectified linear transformation to ensure non-negative firing rates:

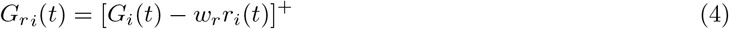

Here, *τ* represents the suppression time constant, *w*_*r*_ is the suppression scaling factor, and [*·*]^+^ denotes the ReLU (rectified linear unit) function. The differential equation was solved using Euler’s method, and the aggregate fMRI signal was derived by averaging *G*_*ri*_(*t*) across the entire cell population.

To implement the nonlinearity hypothesis [22], we first computed the moving average of the activity of grid cell *i, G*_*ci*_(*t*), over a temporal window of *T* seconds:

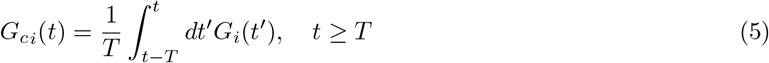

A nonlinear transformation was then applied by raising the averaged activity to the power of *x*, yielding the final activity *G*_*ni*_(*t*):

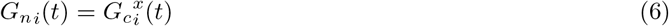

Consistent with our models, the simulated fMRI signals were derived from the population average of all *G*_*ni*_(*t*) values.

### Trajectories

In the star-like trajectory condition, each segment consisted of a single straight path in a single direction, with time *t* reset to zero at the beginning of each segment. All directions were sampled once within one block. Conversely, for the piecewise-linear trajectories, time was reset only at the start of each block. The initial segment began at the designated starting phase, with each subsequent segment originating from the terminal location of the previous one, separated by a pause of *t*_*p*_ seconds. We restricted the turning angles between segments to multiples of 60° every two turns (Fig. 3A). Only segments following these restricted turns were included in the modulation analysis, and angles from 0 to *π* were sampled by these segments. We systematically varied segment lengths to evaluate their influence on the signal, as discussed in the next section.

**Fig 3.**
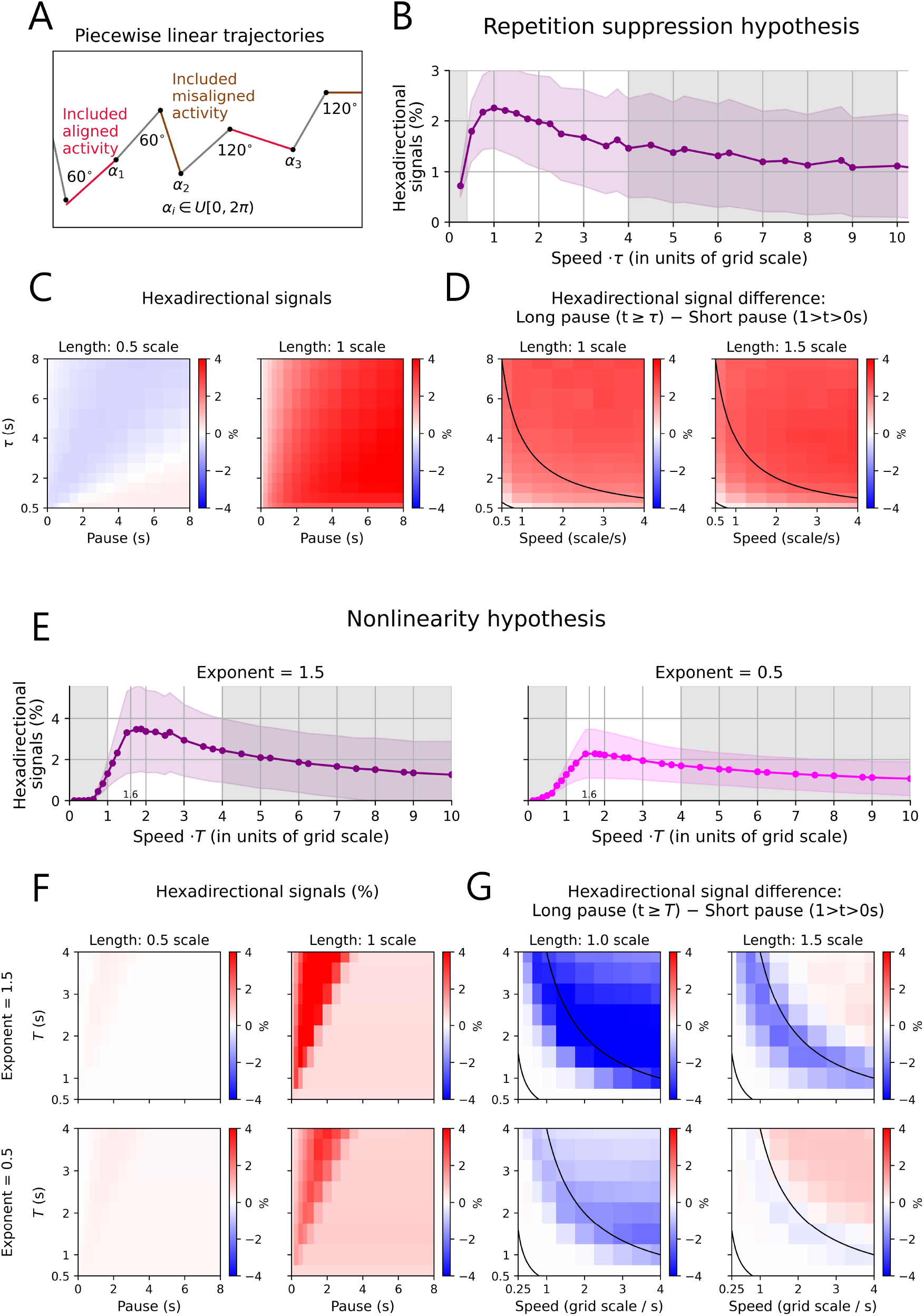
Effects of movement speed, path-segment length, and inter-segment pause duration on hexadirectional signals for piecewise linear trajectories. **A**. Representative piecewise linear trajectory. The calculation of the hexadirectional signal is restricted to the red segments whose onset follows a rotation by a multiple of 60°. **B,E**. For the piecewise linear trajectory, speed has a single-peaked, right-skewed effect on hexadirectional signals under both the repetition-suppression and nonlinearity hypotheses. **C,F**. The hexadirectional signal decreases strongly for short path segments under the repetition-suppression and nonlinearity hypotheses. **D,G**. A longer pause between segments increases the hexadirectional signals under the repetition-suppression hypothesis, but decreases or has minimal effect under the nonlinearity hypothesis.

### Parameters

To ensure that our results were scale-independent, the traveling speed *v* and path length were normalized to units of one grid scale per second and one grid scale, respectively. Consequently, the absolute grid scale does not influence the simulation outcomes. The unit of time was one second.

For the conjunctive grid-by-head-direction cell model, the von Mises concentration parameter was set to *κ* = 8 [21], and the “arm” length of the star-like trajectories was fixed at 2*λ*. We performed a grid search across speeds *v* = 0.25*k* for *k* ∈{1, 2, …, 16}. To maintain consistent spatial sampling across different speeds, the simulation time step *dt* was adjusted to *dt* = 1/10*v*.

For the repetition-suppression hypothesis, the scaling factor was fixed at *w*_*r*_ = 0.5. In the star-like trajectory condition, we defined the “arm” length as max(3, *vτ*) to ensure sufficient time for the suppression dynamics to unfold. We conducted a grid search across speeds *v* ∈ {0.2, 0.4, 0.6, 0.8, 1, 1.2, 1.4,1.6, 1.8, 2, 2.5, 3, 3.5, 4, 5} and suppression time constants *τ* ∈{0.5, 1, 2, 3}. For piecewise-linear trajectories, the parameter space included segment lengths *l* ∈{0.5, 1,1.5, 2, 2.5, 3, 3.5, 4, 5, 6, 7, 8}, speed *v* ∈ {0.5, 1, 1.5, 2, 2.5, 3. 3.5, 4}, time constants *τ∈* {0.5, 1, 1.5, 2, 2.5, 3, 3.5, 4, 5, 6, 7,8}, and pause durations *t*_*p*_ ∈ {0, 0.25, 0.5, 0.75, 1, 1.5, 2, 2.5, 3, 3.5, 4, 5, 6, 7, 8}. To ensure numerical stability and consistent spatial resolution, the simulation time step was dynamically set to *dt* = min(1/10*v, τ* /10).

For the nonlinearity hypothesis, the “arm” length in star-like trajectory simulations was defined as *vT*.

We conducted a grid search across a wide range of parameters, including speeds *v* ∈{0.1, 0.2, 0.4, 0.6, 0.8, 1, 1.2, 1.4, 1.6, 1.8, 2, 2.2, 2.4, 2.6, 2.8, 3, 4, 5}, time windows *T* ∈{1, 2}, and exponents *x∈{*0.5, 1.5 *}*. For the piecewise-linear trajectories, the grid search included segment lengths *l* ∈{0.5, 1, 1.5, 2, 2.5, 3.5, 6}, velocities *v*∈{0.25, 0.5, 0.75, 1, 1.5, 2, 2.5, 3, 3.5, time windows} *T* ∈{0.5, 1, 1.5, 2, 2.5, 3, 3.5, 4}, pause durations *t*_*p*_ ∈{0, 0.25, 0.5, 0.75, 1, 1.5, 2, 2.5, 3, 3.5, 4, 5, 6, 7, 8}, and exponents *x* ∈{0.5, 1.5}. To maintain a consistent spatial resolution relative to the grid scale, the simulation time step was fixed at *dt* = 1/10*v*.

All insets in Figs. 2C and 2D were generated using star-like trajectories. For the insets in Fig 2C, we set *v* ∈{0.2, 0.8, 10} and *τ* = 1. The activity time series in the left inset was simulated using a grid cell with phase zero, with the trajectory originating at phase zero and proceeding in a 10° direction for 10*λ*. We visualized the activity between 3s and 23s, a period during which the virtual agent traversed 4*λ*. For the right inset, the same grid cell and trajectory were used, but the activity was displayed between 0.6s and 1s, again corresponding to a displacement of 4*λ*. For the insets in Fig. 2D, we used *v* ∈{0.2, 1.6, 10} and *T* = 1.

### Quantification of hexadirectional signals

To quantify the hexadirectional signals *H*, we first calculated the average aligned activity 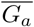, with traveling directions within 15° of the grid axes. The remaining values were averaged to obtain the mean misaligned activity, 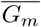. Then, for the repetition-suppression hypothesis and the nonlinearity hypothesis with an exponent smaller than unity, we set 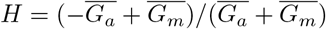. For the other two hypotheses,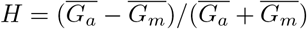.

To quantify the frequency of encountering the same cell in Fig. 2C, we first defined its firing field using a half-maximum threshold; specifically,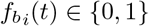 indicates whether the virtual agent was within the firing field of grid cell *i* at time *t*, where a field was defined by *G*_*i*_(*t*) *>* 1/2 max *{G*_*i*_*}*. The unweighted frequency was calculated by averaging 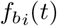 across all time points, cells, and trajectories. For the weighted frequency, we first applied a low-pass filter to the baseline activity *G*_*i*_(*x*(*t*)) using an exponential decay kernel exp(*−t*′/*τ*) with a suppression time constant *τ*. The resulting weight was defined as *G*_*ei*_(*t*) = (*G*_*i*_(*t*′)*exp(*−t*′/*τ*))(*t*) + 10^*−*3^, where the constant 10^*−*3^ ensures non-zero weights. Finally, the weighted frequency was computed as the average of 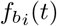 weighted by *G*_*ei*_(*t*) across all dimensions.

To evaluate the variability of time-window averaged grid-cell responses in Fig. 2D, we calculated the standard deviation of the responses, *G*_*c*_(*t*). For each grid cell and trajectory, we first computed the time-window-averaged activity, denoted as 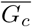. We then calculated the standard deviation of these averaged values 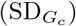 across the entire population of grid cells and all trajectories.

### Statistical tests

To evaluate the effect of speed under the conjunctive-grid-by-head-direction-cell hypothesis, we fitted a linear regression model with hexadirectional signals (*H*) as the dependent variable and speed as the independent variable, testing whether the regression slope significantly deviated from zero. For the hypotheses that depend on grid-like firing patterns, we employed two-sided independent-samples t-tests to assess speed effects. Statistical testing was omitted for star-like trajectory simulations due to the negligible variance observed (see shaded areas in Fig. 2C, D). The influence of trajectory length and pause duration on the modulation strength was assessed using two-sided one-sample t-tests, with p-values corrected for multiple comparisons using the Benjamini–Hochberg False Discovery Rate (FDR) procedure.

## Results

For this study, fMRI signals were simulated by summing the activity of a population of grid cells with varying spatial phases but identical grid orientation and scale *λ*, defined as the distance between the peaks of two neighboring firing fields of a grid cell, as would be the case in a single grid module. We first examined the effect of movement speed on hexadirectional signals. For simplicity, we used trajectories along the twelve “arms” of a star-like figure (Fig. 2A) and assumed that the arms were visited in random order. The length of each arm was fixed. This trajectory configuration has been used experimentally [30].

To simulate the Conjunctive Direction hypothesis, we can confer direction-selectivity onto a random subset of simulated grid cells and single out a periodic set of directions as being preferred by subsets of grid cells. This creates a direct link between cell tuning properties and the population signal. As expected, speed, measured as the number of grid scales traversed per second, did not affect the hexadirectional signals (Slope: 95%, CI [-0.024, 0.018], *p* = 0.778, Fig. 2B). On the other hand, we can make some cells increase their responses weakly with velocity, provided that the velocity aligns with the axes. This is sufficient to support the hexadirectional signal at the population level, with the modulation increasing with speed (Slope: 95%, CI [0.464, 0.506], *p <* 0.001, Fig. 2B).

The other two hypotheses rely on indirect effects mediated by a characteristic time scale. For instance, the repetition-suppression hypothesis invokes a fixed time constant of adaptation (Fig. 2C, see also [21]). Starting from the center of a firing field, on-axis trajectories pass through firing fields every *λ* (the length scale), while off-axis trajectories intersect with firing fields every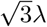. Averaging across all start points, the grid cells on off-axis paths suffer less suppression. There will be an ideal speed *v* for which the difference between on- and off-axis paths is greatest. At low speeds (e.g., *v × τ* = 0.2*λ*, Fig. 2C, left inset), a grid cell almost completely recovers from the field-mediated suppression before another grid field is encountered, regardless of the travel direction; as a consequence, the activity difference between aligned and misaligned directions vanishes. At high speeds (e.g., *v×τ* = 10*λ*, Fig. 2C, right inset), the fast movement effectively averages activity over time. Only at intermediate speeds (e.g., *v×τ* = 0.8) will hexadirectional signals emerge.

Neurovascular coupling is presumed to be nonlinear, making the fMRI signal sensitive not only to the mean, but also to the variance of the firing rate. The nonlinearity hypothesis assumes a constant time window *T* (Fig. 2D), over which the activities are averaged. After that, a nonlinearity is applied, representing the neurovascular coupling. Similar to the phenomena seen under the repetition-suppression hypothesis, the amplitude of the hexadirectional signal exhibits a bell-shaped dependence on speed. At low speeds (e.g., *v×T* = 0.2*λ*, Fig. 2D, left inset), the hexadirectional modulation is negligible, as the activity averaged over *T* is uniformly variable across movement directions. At high speeds (e.g., *v×T* = 10*λ*, Fig. 2D, right inset), the variance across different trajectories is high only along the exactly aligned direction. High speeds lead to long paths that cross multiple grid fields; along paths exactly aligned with the lattice, the same spatial phase is reached after traversing any integer multiple of the lattice scale. In such a case, the difference between different paths does not diminish with longer paths, which implies that the variance across paths remains high. On the other hand, in other directions, the path veers off the restricted set of spatial phases characteristic of the lattice axes; with more spatial phases sampled on such paths, the variance of different paths diminishes. Consequently, the hexadirectional signal is maximal at moderate speeds (e.g., *v×T* = 1.6*λ*).

To determine whether the single-peaked speed effect generalizes to more complex paths, we next employed a piecewise-linear trajectory (Fig. 3A). In this simulation, we systematically varied both the length of the linear path segments and the pause duration between such segments. While these manipulations increased the overall variance of the results, the core finding remained robust. As expected, we observed a similar unimodal speed effect on the hexadirectional signals (Fig. 3B, E). Under the repetition-suppression hypothesis, the activity at *v×τ* = 1 *λ* was higher than that at *v ×τ* = 0.25*λ* (t(71065.93) = 400.62, *p <* 0.001, CI95% = [1.52, 1.54], Cohen’s d = 2.20) and *v× τ* = 10*λ* (t(107998) = 204.47, *p <* 0.001, CI95% = [1.13, 1.15], Cohen’s d = 1.24). Under the nonlinearity hypothesis with exponent 0.5, the activity at *v× T* = 1.5*λ* was higher than that at *v×T* = 0.25 (t(52559.04) = 432.55, *p <* 0.001, CI95% = [2.23, 2.25], Cohen’s d = 2.23) and *v×T* = 10*λ* (t(55002.47) = 157.13, *p* < 0.001, CI95% = [1.20, 1.23], Cohen’s d = 1.10). Similar results were observed with an exponent of 1.5 (comparing *v×T* = 1.5*λ* with *v× T* = 0.25*λ*: t(52499.00) = 379.36, *p <* 0.001, CI95% = [3.30, 3.33], Cohen’s d = 1.96; and comparing *v×T* = 1.5*λ* with *v×T* = 10*λ*: t(47230.72) = 144.31, *p <* 0.001, CI95% = [2.02, 2.08], Cohen’s d = 1.08).

Second, the effect of the path-segment length distinguishes the two conjunctive-cell hypotheses from the two hypotheses that depend on grid-like-firing patterns. Under the conjunctive-cell hypotheses, hexadirectional signals emerge regardless of segment length. Conversely, hexadirectional modulation only appears for longer path segments under the other two hypotheses (Fig. 3C, F). In particular, under the repetition-suppression hypothesis, reducing the length from *λ* to *λ*/2 strongly decreased the hexadirectional signal. At a length of *λ*/2, only 13.9% of parameter combinations showed significant positive hexadirectional modulation (*t*-test per combination across 100 simulations, Bonferroni-corrected *p <* 0.05; *M* =*−*0.321 ± 0.291 [mean ± SD], range [ *−*0.663, 0.282]), whereas at a length of one *λ*, 85.6% were significantly positive (*M* = 2.540 ± 1.107, range [0.125, 4.164]). We observed similar results under the nonlinearity hypothesis (Fig. 3F). Pooling across both exponent values, significantly positive values dropped from 85.6% at a length of one *λ* (*M* = *−*1.310 ± 1.295, range [0.312, 6.318]) to just 1.7% at a length of *λ*/2 (*M* = 0.090 ± 0.083, range [ −0.103, 0.314]). Under the two hypotheses that depend on grid-like-firing patterns, this phenomenon might thus provide a way to measure the spatial scale of grid-cell lattices non-invasively.

Finally, to further differentiate between the repetition-suppression and the nonlinearity hypotheses, we examined the effect of pauses between trajectory segments connected by turns that followed specific rules: Every second turn was a multiple of 60°, the other turns were chosen randomly. Only segments following the 60° turns were used to measure the hexadirectional modulation. Each turn was associated with a pause that ranged from short (less than a second) to longer than the critical time scale (the suppression time constant *τ* or the time window *T* from the nonlinearity hypothesis). Under the repetition-suppression hypothesis, each combination of starting phase and movement direction engages and suppresses a specific set of cells. A turn changes the engaged cell population and is therefore disruptive. Increasing the pause between segments allows the suppression to (partially) recover, reducing the disruptive effect of the turn. In contrast, under the nonlinearity hypothesis, a 60° turn is beneficial, particularly for short segments.

Increasing the pause weakens this benefit and instead introduces additional disruption. For path segments with lengths between 1.0*λ* −1.5 λ and speeds within the central, unshaded regions in Figs. 3B, E, active repetition-suppression meant that longer pauses significantly enhanced the hexadirectional signals (Fig. 3D; *M* = 1.654 ± SD = 0.625, range [0.295, 2.552]). In contrast, the longer pauses in the nonlinear model failed to increase the modulation strength (Fig. 3G; *M* = −0.633 ± SD = 1.037, range [ *−*4.725, 0.108]). The contrast between the effects under the two hypotheses persists even for longer segment lengths, albeit less prominently (see Supplementary Figs. 1–3).

If multiple mechanisms play a role in causing the hexadirectional signals, the correspondence between experimental findings and underlying hypotheses becomes complex (Table 1). If the signals fail to increase with longer segments, hypotheses that depend on grid-like-firing patterns are ruled out. Moreover, if hexadirectional signals effectively vanish for short path segments, the conjunctive-cell hypotheses are ruled out. For this reason, the path-segment length is the most informative manipulation for distinguishing between these two classes of hypotheses. Other features support particular theories, but do not rule out the presence of other mechanisms. For instance, a peak in the modulation strength versus speed can coexist with some neurons being sensitive to specific movement directions or velocity vectors. Finally, varying the pause duration might produce no measurable effect under three of the hypotheses, so we cannot exclude the possibility that these mechanisms are present. Indeed, the possibly negative effect of pauses under the nonlinearity hypothesis might counteract the pause-induced release from the repetition-suppression hypothesis. Consequently, the net outcome might remain undetermined.

**Table 1.** Possible experimental findings and their interpretations. Assuming that the four hypotheses discussed here represent the only possible mechanisms underlying the hexadirectional modulation of fMRI signals, the symbols indicate the following interpretations: ✓: at least one of these checked hypotheses is supported; **?**: the hypothesis may be either true or false; ***×***: the hypothesis is incompatible with the experimental findings.

| Experimental Finding | Direction-selective-cell hypothesis | Velocity-selective-cell hypothesis | Repetition suppression hypothesis | Nonlinearity hypothesis |
| --- | --- | --- | --- | --- |
| **Effect of speed** |  |  |  |  |
| Positive | ? | ✓ | ? | ? |
| Single-peaked | ? | ? | ✓ | ✓ |
| Null | ✓ | × | × | × |
| **Does the hexadirectional signal disappear with short path segments?** |  |  |  |  |
| Yes | × | × | ✓ | ✓ |
| No | ✓ | ✓ | ? | ? |
| **Does the hexadirectional signal increase with longer path segments?** |  |  |  |  |
| Yes | ? | ? | ✓ | ✓ |
| No | ✓ | ✓ | × | × |
| **Effect of pause duration** |  |  |  |  |
| Positive | ? | ? | ✓ | ? |
| Null | ✓ | ✓ | ? | ✓ |
| Negative | ? | ? | ? | ✓ |

## Discussion

While grid-cell activity is believed to underlie hexadirectional modulation of fMRI signals, the exact mechanism linking the two remains unclear. Four hypotheses have been proposed: conjunctive direction-selective grid cells [20–22], conjunctive velocity-selective grid cells, repetition suppression [20, 21], and nonlinear transformations of single-cell activity into fMRI signals [22]. However, none of these hypotheses has received strong experimental support. In the current study, we, therefore, developed experimentally testable approaches to distinguish among the four hypotheses.

If grid-like-firing patterns are the source of hexadirectional modulation and either of the proposed grid-cell-based hypotheses holds true, then we have found a way to noninvasively estimate the grid scale by varying the path length and measuring the hexadirectional effect. Indeed, the hexadirectional signal should effectively disappear when the subjects only travel along a short path, that is, a path shorter than half the grid scale, and become strongest when the path length is about one grid scale.

Movement speed links the spatial and temporal scales of grid-cell firing, making speed a marker for the origin of hexadirectional modulation in the fMRI signal. There could be an ideal speed that maximizes the modulatory effect, which in itself would lend evidence to the two hypotheses that require grid-like firing patterns.

Grid cells are organized into multiple modules that differ in grid scale and may also exhibit distinct grid orientations [31]. Hexadirectional analyses should, therefore, not be conducted using signals averaged over larger brain regions. Thanks to the partial anatomical segregation of the grid modules, hexadirectional analyses can, however, be carried out for smaller, spatially localized voxel clusters.

Neither of the conjunctive tuning hypotheses will have these particular scale- and speed-dependent effects. Pauses between travel periods, moreover, will differentially affect the fMRI signal under the competing theories: Longer pauses increase the hexadirectional signal under repetition suppression, but tend to diminish the modulation under the nonlinearity hypothesis.

Therefore, we propose the following road map for studying how hexadirectional signals arise: first, test the effect of path segment length; secondly, vary the speed of movement; thirdly, introduce pauses between travel segments of varying duration.

We do argue that the standard explanation [20] for the repetition-suppression hypothesis is incomplete.

Critically, when averaged across all starting positions, the mean frequency of encountering the same cell does not vary by movement direction (Fig. 2C, orange lines). The expected difference between aligned and misaligned directions appears, however, when we weigh the average by recent activity (Fig. 2C, black lines). Interestingly, this weighted-average frequency also slightly increases in misaligned directions, despite those directions experiencing less suppression. This nuance highlights a gap in our understanding of the repetition-suppression hypothesis, calling for a detailed analysis beyond the scope of this study. Although the biological basis for the nonlinearity hypothesis is still unclear, a number of nonlinearities exist in the transformation of the single-cell activity to fMRI signals [26–29, 32–35].

Our results are consistent with other studies [21, 22, 25] that investigated the feasibility of the mechanisms to generate hexadirectional signals, while we here have focused on proposing ways to examine them empirically. Bin Khalid et al.’s extensive simulations indicate that for realistic parameters, direction-selective cells and repetition suppression cells can produce a hexadirectional modulation, with direction-selective cells having a stronger effect. Bin Khalid et al. introduced a structural-functional-mapping hypothesis, too, but the effects are small, so we do not include this hypothesis in the current study. Almog et al. studied the effect of a power-law nonlinearity, with the result that hexadirectional signals only emerge for exponents larger than unity. Almog et al. treats speed as a constant, calculating a single cell’s average activity over a fixed path length before applying the nonlinearity. In the present study—where speed is variable—we have opted for an average over a fixed time window. Both Almog et al. and [25] find that, at least in the rodent, the preferred directions of rodent direction-selective cells do not align with the grid axes. Such meta-analyses of rodent grid-cell data from several laboratories do not rule out a role for direction-selective cells in human beings, so we included this hypothesis in the current study.

Simulations predict stronger hexadirectional modulation of population activity than observed in fMRI experiments. There are several reasons why this might be the case. First, our simulations consist entirely of grid cells, whereas an fMRI voxel likely contains a heterogeneous population of neurons, which reduces the signal-to-noise ratio. Second, biological grid cells rarely exhibit the idealized firing patterns used in this modeling study. Third, the parameters governing signal magnitude—such as the suppression scaling factor (*w*_*r*_ = 0.5 in our model)—may be smaller *in vivo*. Our predictions, therefore, concern relative effect sizes of different manipulations, not absolute values. Some researchers do not find hexadirectional fMRI signals at all [36, 37], suggesting that the effect may be subtle and highly sensitive to experimental protocols.

Moreover, apparent hexadirectional signals may arise from analysis procedures that presuppose stochastic uniformity that is not present in the data. In fact, Almog et al. argue that, without careful analysis, signals from a population of direction-selective cells may be misclassified as hexadirectional [22]. Such cells are abundant in the entorhinal cortex [38, 39]. Under this account, previously reported correlations between hexadirectional signals and behavioral performance [10, 16, 20] may instead be explained by the tuning properties of direction-selective cells.

## Acknowledgments

We are grateful to all members of the Flanagin and Herz labs for their input and thank Richard Kempter for helpful comments on the manuscript.

## Supporting information

**S1 Fig.**
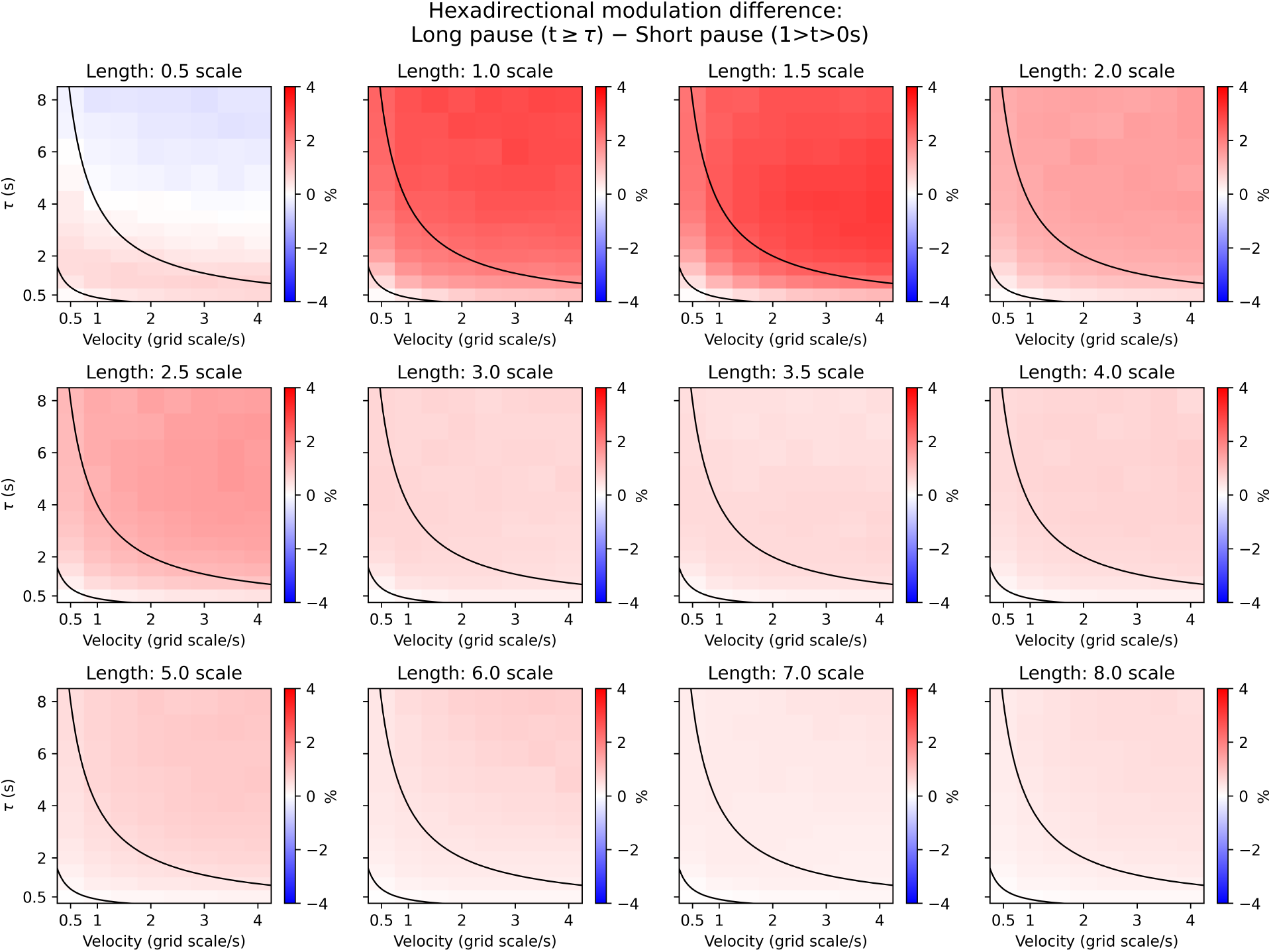
Persistence of the pause effect with even longer segment lengths under the repetition suppression hypothesis. Simulation results for the differences in hexadirectional modulation between short and long pauses are shown as a function of velocity and path segment length, and for different suppression time constants.

**S2 Fig.**
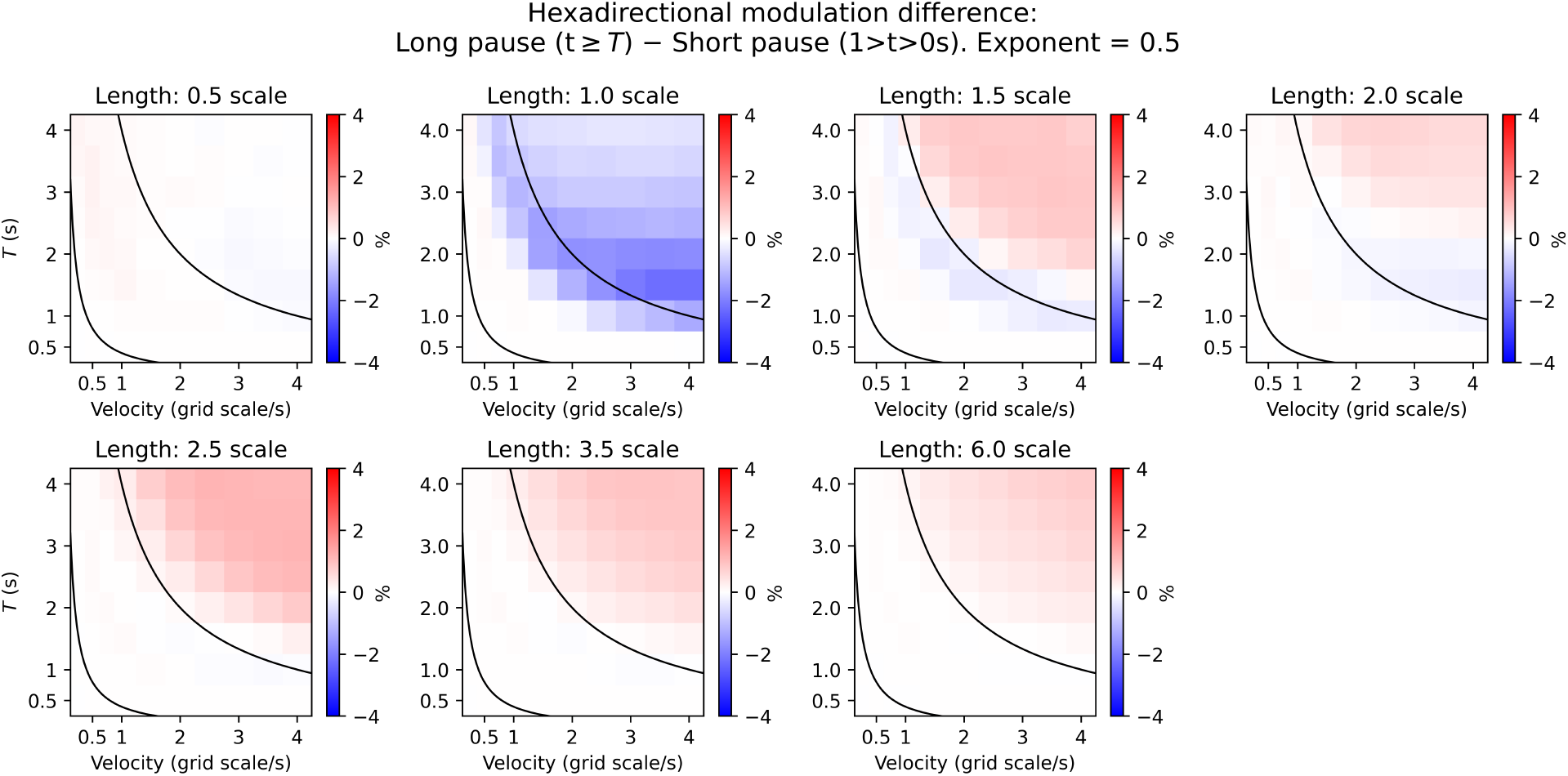
Effect of pauses under the nonlinearity hypothesis with an exponent of 0.5. As in S1 Fig., but under the nonlinearity hypothesis and for different nonlinearity time windows.

**S3 Fig.**
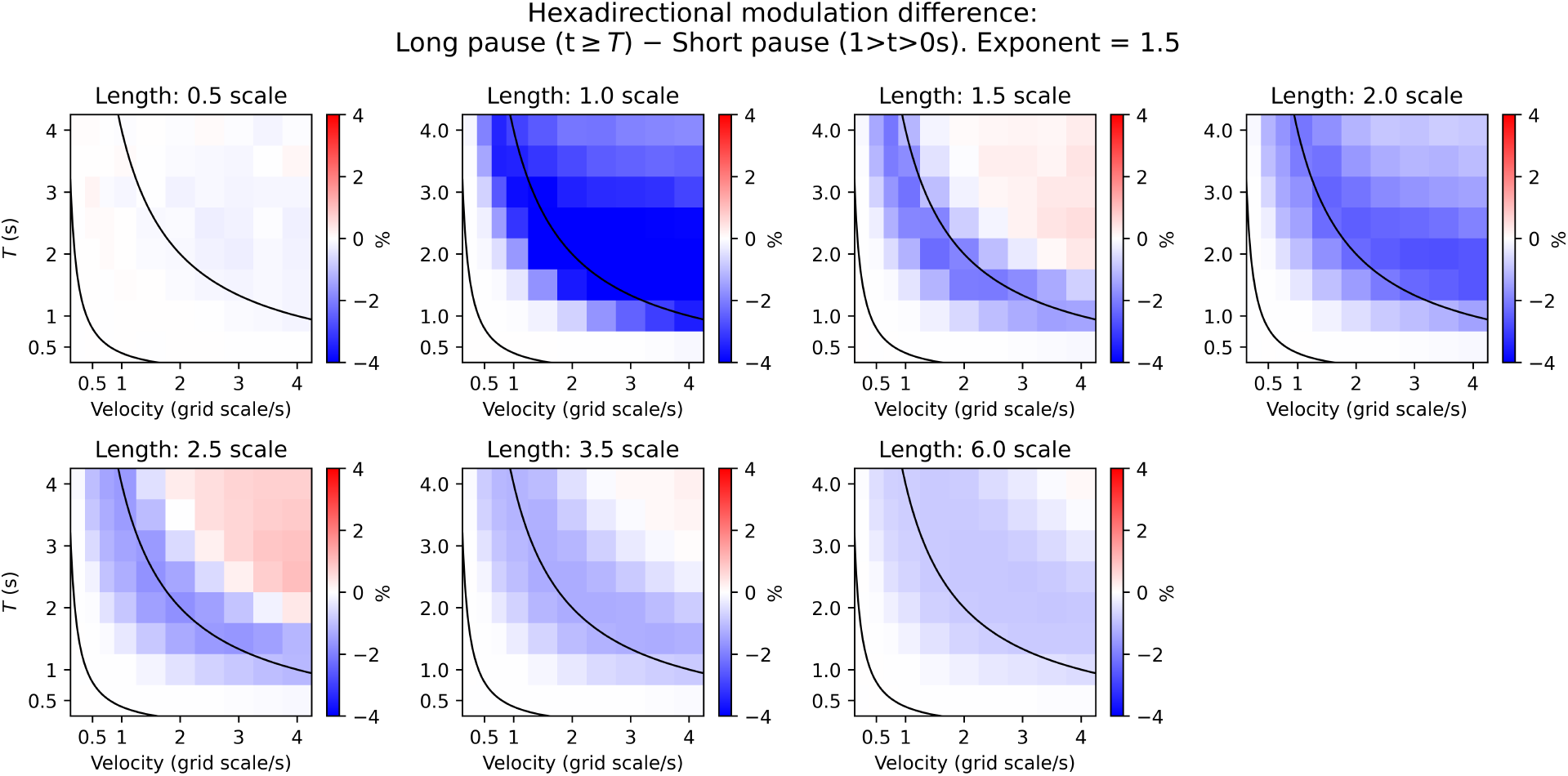
Effect of pauses under the nonlinearity hypothesis with an exponent of 1.5. Same as in S2 Fig. The effect is consistent, albeit weaker, with longer path segments.

